# Real-time tracking of Tomato brown rugose fruit virus (ToBRFV) outbreaks in the Netherlands using Nextstrain

**DOI:** 10.1101/2020.06.02.129395

**Authors:** Bart T.L.H. van de Vossenberg, Michael Visser, Maaike Bruinsma, Harrie M.S. Koenraadt, Marcel Westenberg, Marleen Botermans

## Abstract

Tomato brown rugose fruit virus (ToBRFV) is a Tobamovirus that was first observed in 2014 and 2015 on tomato plants in Israel and Jordan causing discolorations and deformation of leaves and fruits. Apart from tomato, damage in pepper fruits has been reported. Since the first description, the virus has been found on all continents except Oceania and Antarctica.

In October 2019, the Dutch National Plant Protection Organization received a tomato sample suspected to be infected with ToBRFV as part of an official specific survey of tomato fruit growers in the Netherlands carried out July to October 2019. During the survey 124 companies were visited and inspected for possible ToBRFV symptoms. Of the 47 samples tested, one sample tested positive with ELISA and test plants, which was verified using real-time RT-PCR and Illumina RNAseq data. A follow-up survey was initiated to determine the extent of ToBRFV presence in the Dutch tomato horticulture and identify possible linkages between ToBRFV genotypes, companies and epidemiological traits. We used Nextstrain to visualize these potential connections.

Genomic diversity of ToBRFV isolates found in the Netherlands group in three main clusters which are hypothesized to represent three sources. No correlation was found between genotypes, companies and epidemiological traits such as rootstock and scion varieties, seed batches and nurseries, and the source(s) of the Dutch outbreak remain unknown.

This paper describes a Nextstrain build containing ToBRFV genomes up to and including November 2019. The NPPO-NL has committed itself to maintain and improve the build. Sharing data with this interactive online tool will enable the regulatory plant virology field to better understand and communicate the diversity and spread of this new virus. Other organizations are stimulated to share data or materials for inclusion in the Nextstrain build, which can be accessed at //40.91.255.14/ToBRFV/20191231.

## Introduction

Tomato brown rugose fruit virus (ToBRFV) belongs to the genus *Tobamovirus* and was first described by Salem *et al*. in 2016 after a finding in tomato (*Solanum lycopersicum*) plants in Jordan [2]. This appeared to be the same virus that was causing problems since 2014 in Israel on tomato plants showing mosaic patterns on leaves, and occasionally narrowing of leaves and yellow spotted fruits. The tomato *Tm-2^2^ R*-gene, which confers resistance to Tobacco mosaic virus (TMV) and Tomato mosaic virus (ToMV), offers no protection against ToBRFV. Hence the virus is regarded a major threat for tomato crops (1, 2). Pepper crops (*Capsicum* spp.) can also be affected by the virus, producing symptoms similar to those observed on tomato (3). However, outbreaks in pepper have only been reported from Mexico, Jordan and Italy (3–5). Several other plant species, including *Petunia* spp. and *Solanum nigrum* could be inoculated under laboratory conditions (1). As with other tobamoviruses, ToBRFV is stable and can easily be transmitted by contact (e.g. farming equipment, hands, clothing and plant-to-plant contact) (6), and propagation material (grafts, cuttings). Seed transmission of ToBRFV is presumed (7), and bumblebees have been reported to serve as a vector for ToBRFV and potentially contribute to the spread of the disease (8).

ToBRFV has a similar genome architecture compared to other tobamoviruses, and consists of a ~6.4 kb monopartite linear single stranded RNA (ssRNA) encoding four proteins: the small replicase subunit, a RNA-dependent RNA polymerase (RdRp) which is translated through a suppression of translation at the end of ORF1, a movement protein (MP), and a coat protein (CP) (2). To date, nine complete ToBRFV genome sequences have been deposited in NCBI Genbank (MK133095, MN167466, MK319944, KX619418, MK165457, MN182533, MK648157, MK133093, KT383474) representing outbreak locations in Middle East, North-America and Europe.

Following the initial findings in Jordan and Israel, the virus was reported from the State of Palestine (9), Mexico (10), the United States of America (11), Germany (12), and Italy (13) in 2018, and Turkey (14), China (15), the United Kingdom (16), the Netherlands (17), Greece (18), and Spain (19) in 2019. In 2020, ToBRFV outbreaks were reported from France (20) and Egypt (21). Early October 2019, the National Reference Center for plant health of the Dutch National Plant Protection Organization (NPPO-NL) received a tomato sample as part of an official specific survey of tomato fruit growers, that was suspected to be infected with ToBRFV. Testing of the sample by DAS-ELISA, bio-assay and real-time RT-PCR (22) further strengthened the initial suspicion. The sample was subjected to Illumina RNAseq analysis for final confirmation. Comparison of the complete viral genomes present in the sample with public NCBI accessions verified the presence of ToBRFV. ToBRFV is regulated as part of EU Commission implementing decision 2019/1615 as of 1 November of 2019 (23). Therefore the initial finding triggered a follow-up survey targeting the nursery and the seed lots of the affected plants as well as intensified specific surveillance by the NPPO-NL to determine the extent of ToBRFV presence in the Dutch tomato horticulture. This resulted in new ToBRFV findings at tomato fruit companies and the enforcement of measures to prevent spread of the virus.

The aim of the current study was to determine the genomic diversity that was represented within and between the Dutch outbreak locations to identify possible linkages with epidemiological factors such as scion and rootstock varieties. We used Nextstrain (24), a collection of open-source bio-informatic tools, to create an interactive view of the diversity and spread of the virus. Real-time virus genome sequencing and rapid dissemination of epidemiologically relevant results, which is central in the Nextstrain philosophy, has been mainly applied to viruses causing disease in humans, such as Zika and Ebola (25). By sharing the Dutch outbreak sequences in the context of geography and epidemiological traits at an early stage, we believe this will empower NPPOs and the research community to better understand and battle this important viral disease that threatens tomato and pepper production worldwide.

At the time of preparation of this publication, public ToBRFV sequences and sequences generated in this study dating from November 2014 to November 2019 were included in the ToBRFV Nextstrain build. The Dutch NPPO has committed itself to keep adding sequence data and improving the build. This paper also serves as an invitation to other NPPOs to contribute their data to improve the predictive power of the tool. The ToBRFV Nextstrain build can be accessed via //40.91.255.14/ToBRFV/20191231 (last accessed 13 May 2020).

## Material and methods

### Sampling

Following the initial ToBRFV finding, tomato greenhouses were selected based on an assessment including potential linkages to the initial outbreak location and data collected from private laboratories, and were inspected by phytosanitary inspectors of NPPO-NL. Typically, 50 young leaf parts were sampled from symptomatic tomato plants per greenhouse. When no symptomatic plant parts were found, asymptomatic material was sampled. Per company, one or more samples consisting of 50 young leaf parts were taken. In cases where there was a strong suspicion of ToBRFV but no young leaf parts available, additional material was sampled such as tomato roots and/or water from rock wool. In addition to the survey samples taken from Dutch greenhouses, samples from regular import inspections suspected to contain ToBRFV were included in this study.

Leaf samples were split into two subsamples of 25 leaves each. From each subsample, a leaf disc was obtained using a 6 mm diameter leaf punch. Subsample punches were added to 5 mL GH+ buffer (26) (6 M guanidine hydrochloride, 0.2 M sodium acetate pH 5.2, 25 mM EDTA, and 2.5% PVP-10) spiked with Bacopa chlorosis virus (BaCV) to a Cq value in the range of 25-30, which is used as an RNA extraction efficiency control. Punch outs were homogenized with a Homex 6 (Bioreba, Switzerland) prior to RNA extraction with the Sbeadex Maxi Plant Kit (LCG genomics, United Kingdom) on the automated KingFisher Flex 96 platform (ThermoFisher, MA, USA).

Extracted RNA was tested directly, or stored at −20 °C until use. Real-time RT-PCR reactions were performed based on the ISHI-VEG protocol for ToBRFV detection in tomato and pepper seeds (22). Reaction mixes contained two primer-probe pairs targeting the ToBRFV genome, and one targeting the BaCV spike-in (Table 1). Reaction mixes consisted 1x UltraPlex 1-Step ToughMix (Quanta Biosciences, MD, USA), 300 nM of each forward and reverse primer, 200 nM of each probe, and 3 μL of RNA template. Molecular grade water was added to reach a final volume of 25 μL. Real-time RT-PCR reactions were performed in a CFX96 thermal cycler (BioRad, CA, USA) using the following conditions: 50°C for 10 min, 95°C for 3 min, followed by 40 cycles of 95°C for 10 sec and 60°C for 1 min and plate read at 60°C. When one or both ToBRFV primer-probe combinations produced a Cq value of ≤30, the virus was regarded to be detected. When both ToBRFV combinations failed to produce a Cq value ≤30 and the spike-in resulted in a Cq value of ≤32, the virus was not detected. When both ToBRFV combinations resulted in Cq values >30, and the spike-in resulted in a Cq value of >32, the result was undetermined, and testing was repeated.

**Table 1.**
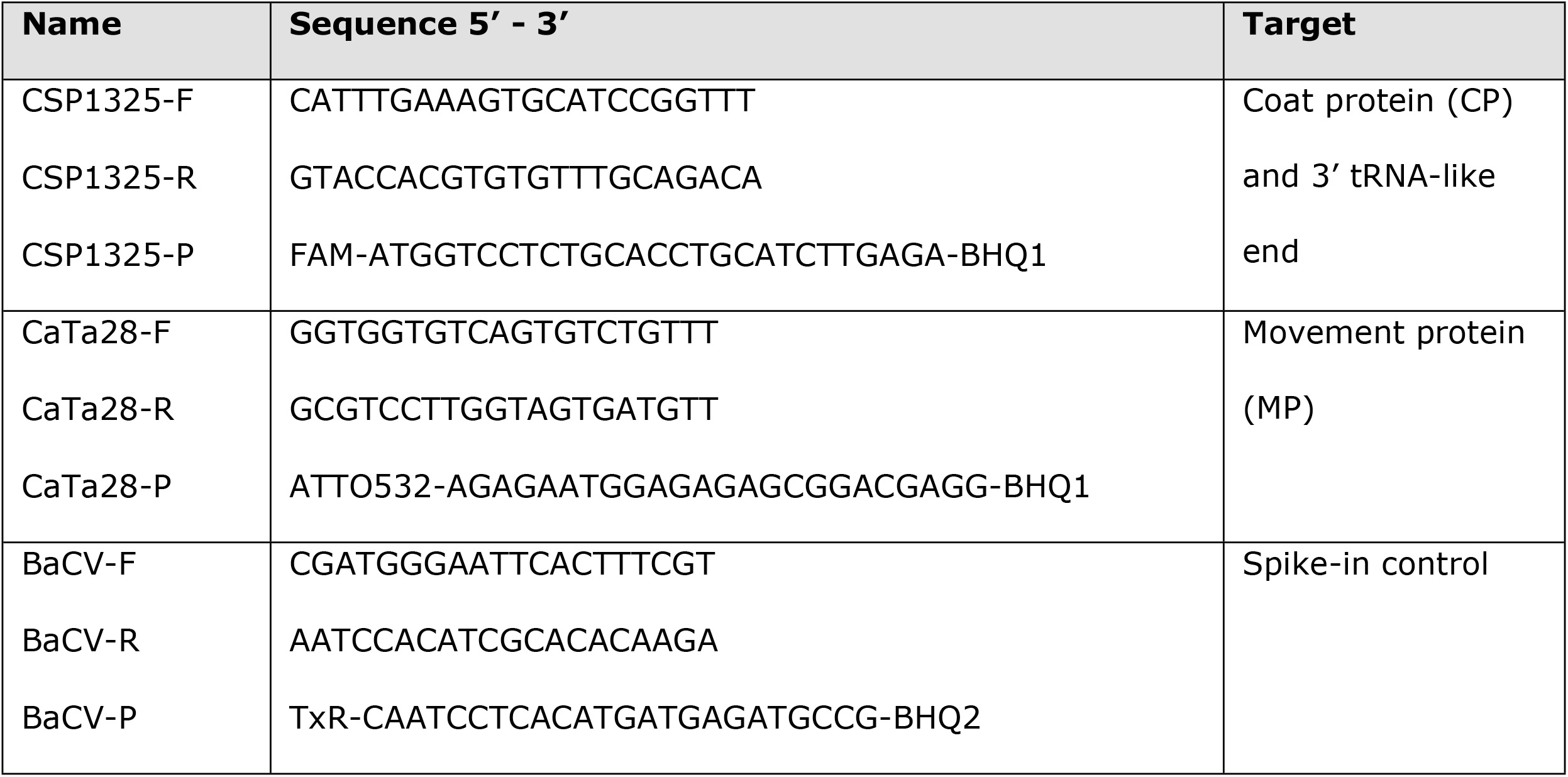
Oligonucleotides used in this study.

### Illumina sequencing

RNA extracts in which the virus was detected were DNAse I treated (Qiagen, Germany) for 15 min at room temperature prior to sequencing. Samples were sent to GenomeScan (Leiden, the Netherlands) for generation of 2 Gb Illumina RNAseq 150PE (paired-end) data per sample. The Ultra II Directional RNA Library Prep Kit for Illumina (New England Biolabs, MA, USA) was used to process the samples according to the protocol “NEBNext Ultra II Directional RNA Library Prep Kit for Illumina”. Briefly, rRNA was depleted from total RNA using the rRNA removal kit Ribo-Zero Plant (Illumina, CA, USA). After fragmentation of the rRNA reduced RNA, a cDNA synthesis was performed which was used for ligation with the sequencing adapters and PCR amplification of the resulting product. Quality and yield after sample preparation were measured with a Fragment Analyzer (Agilent, CA, USA) prior to pooling for sequencing on an Illumina NovaSeq (Illumina, CA, USA).

### Verification pipeline

RNAseq data were uploaded into CLC Genomics workbench v11.0.1 (Qiagen, Germany) and run in a custom workflow build for detection of *de novo* assembled viral contigs (S1 Fig). First, a quality trim (quality limit = 0.05; ambiguous limit = 2) was performed, followed by a *de novo* assembly (map reads back to contigs = on; length fraction = 0.8; similarity fraction = 0.8, minimum contig length = 200) and consensus sequences extraction (low coverage threshold= 10; remove regions with low coverage = on; post-remove action = join). The *de novo* assembled contigs were analyzed using the Basic Local Alignment Search Tool (BLAST, maximum alignments per database sequence = 5; maximum E-value = 1e-6, minimum identity = 70%) with a local installation of the NCBI nr/nt database (27). Blast results were split into plant and non-plant hits, which were visualized in Krona (bit score threshold = 25) (28) (S2 Fig). The same pipeline was repeated with 1% of all reads as *de novo* assembly of high coverage contigs can be problematic resulting in fragmented assemblies. CLC external applications that were used in the pipeline are deposited on GitHub (https://github.com/NPPO-NL/CLC-External-Applications). Putative ToBRFV sequences were manually extracted from the assemblies, and reliability of the assembled putative ToBRFV contigs were determined by visually inspecting read mappings. Consensus sequences were annotated in Geneious R11 (Biomatters, New Zealand) based on the publicly available ToBRFV Tom1-Jo genome KT383474 (2)(minimal sequence similarity = 90%) and the Geneious ORF finder tool (genetic code = table 1, minimum size = 200), and submitted in NCBI GenBank under accessions MN882011 to MN882064. The corresponding SRAs were submitted under accessions ERS4296142 to ERS4296201.

Frequency of major genotypes in the individual samples was determined with the “Basic Variant Calling” tool in CLC Genomics Worbench v11.0.1. (ploidy = 0, minimum coverage = 20, minimum count = 2, minimum frequency = 10%, base quality filter = default) using read-mappings to ToBRFV genome KX619418.

A phylogenetic tree was constructed with the ToBRFV genome sequences generated in this study, together with publicly available ToBRFV sequences. Models for nucleotide substitution of the MAFFT (29) aligned sequences were obtained using the “model testing” tool in CLC Genomics workbench v11.0.1 which was run with default settings for the Hierarchical likelihood ratio test (hLRT), Bayesian information criterion (BIC), Minimum theoretical information criterion (AIC), and Minimum corrected theoretical information criterion (AICc). The clustering analysis was performed with MrBayes v3.2.6 (30) implemented in Geneious R11 under the GTR model with gamma distribution and estimation of invariable sites (GTR + G + I) with a random starting tree and four Monte Carlo Markov Chains for 10^6^ generations. Trees were sampled every 200 generations, and the first 10^5^ generations were discarded as burn-in. Remaining trees were combined to generate a 50% majority rule consensus tree with posterior probabilities.

### Nextstrain implementation

Nextstrain is a bioinformatics pipeline that uses two tools, Augur and Auspice (24). Augur (github.com/Nextstrain/augur) is a bioinformatics toolkit for phylogenetic analysis that uses a series of modules that produces output that subsequently can be visualized by Auspice (github.com/Nextstrain/auspice). Using the augur modules, ToBRFV sequences were aligned with MAFFT (29) and a RAxML clustering (31) was performed. The initial tree was refined with metadata and a time tree was build using TreeTime (32) with optimization of scalar coalescent time. Internal nodes were assigned to their marginally most likely dates, and confidence intervals for node dates were estimated. Amino acid changes were determined based on the functionally annotated Tom1-Jo genome (KT383474 = NC_028478). Additional features had to be added to the NCBI Genbank file (.gb) to enable the use of this sequence as reference, i.e. gene names and translation tables for the four CDS annotations. Next, Augur output was exported and visualized in auspice. The ToBRFV Nextstrain build is deposited on GitHub (https://github.com/NPPO-NL/nextstrain-ToBRFV).

## Results

### Sampling and ToBRFV detection

Following the initial specific survey carried out amongst tomato fruit growers from July-October 2019, by November 2019, additionally greenhouses of 68 companies were inspected and symptoms were occasionally observed, especially young leaves showing chlorotic mottle, mosaic or leaf narrowing (Fig 1). In three cases growers reported delayed ripening or brown necrotic spots on green fruits. From the visited greenhouses a total of 135 samples were taken and split in two or more subsamples that were individually analyzed with real-time RT-PCR. In addition, two fruit samples (39070022 and 39070030) from regular import inspections (which were split into four subsamples) were included for testing. Of these, 70 subsamples representing 21 companies and the two intercepted samples produced positive real-time RT-PCR results in one or both assays with Cq values ranging from 5.0 to 30.1 in the CP assay and from 4.6 to 31.2 in the MP assay. An overview of samples that tested positive with the real-time RT-PCRs is provided in S1 Table.

**Figure 1.**
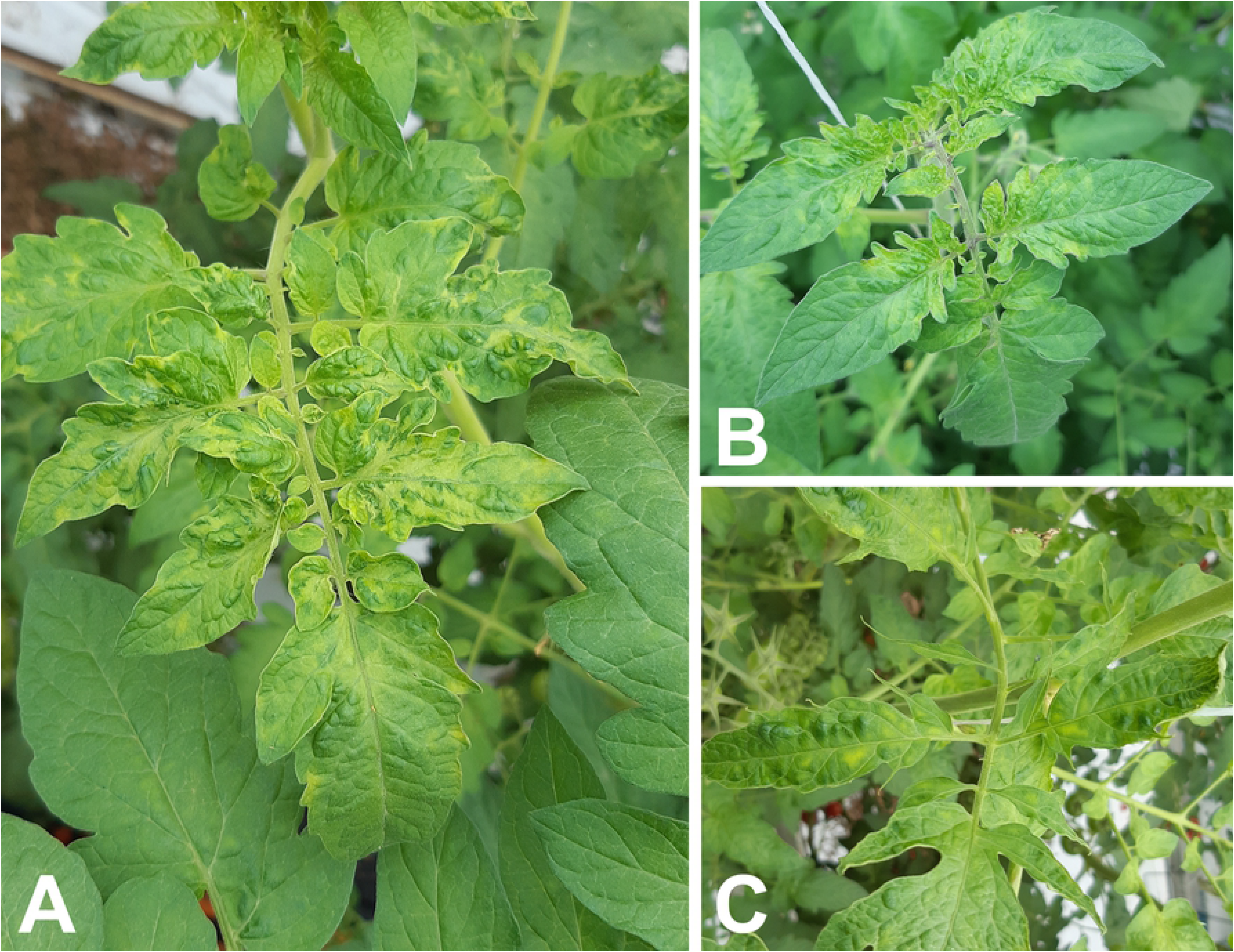
Symptoms observed during phytosanitary inspections. **A** Chlorotic mosaic on young leaves of ToBRFV and PepMV infected tomato plants; **B** Chlorotic mottle on leaf bases of young leaves; and **C** narrowing of and chlorotic mottle on young leaves

### ToBRFV verification using Illumina RNAseq

Plant rRNA depleted RNAseq Illumina data were generated for 70 RNA extracts representing 35 samples taken at phytosanitary inspections of 21 companies growing tomato (or in two cases had been growing tomatoes in the past) in the Netherlands, and 2 samples taken at regular import inspections of tomato fruits. On average 3.7 Gb of sequence data was generated per sample, and *de novo* assembly resulted in an average of 19,645 scaffolds per dataset when using all data. When the analysis with all data failed to produce a single contiguous sequence on account of too many reads that were used in the assembly, this was achieved with the 1% sampled analysis. Contigs producing blast hits for ToBRFV were obtained from 52 datasets representing 17 companies. Visual inspection of the *de novo* assemblies and structural annotation of these contigs verified the identity of the virus in all 52 cases. In two datasets obtained from sample 39941641, two different genotypes were assembled resulting in a total of 54 assembled and verified ToBRFV genomes. The obtained genome size ranged from 6,301 to 6,397 nt with average read coverage ranging from 13 to 579,611x (median coverage = 54,330x). Subsample 39941596_B was sequenced twice (once with and once without DNAse I treatment) in independent sequencing runs as control. The obtained ToBRFV genome sequences were identical, and both sequences represent a unique genotype in the overall dataset.

Subsamples in which ToBRFV could be verified typically had Cq values for the CP and MP assays lower than 25. Only in two cases the complete ToBRFV genome could be constructed from samples with Cq values higher than 25. In some of the samples with Cq values higher than 25, low numbers of reads mapping to the ToBRFV could be obtained, but these were insufficient to verify the presence of ToBRFV in the sample.

Apart from ToBRFV hits, RNAseq data obtained from tomato plants all produced Pepino mosaic virus (PepMV) hits.

### Genomic variation

The obtained ToBRFV sequences were highly similar with 99.3% to 100% identical sites (up to 43 single nucleotide polymorphisms: SNPs). For the 160 amino acid (aa) coat protein, no non-synonymous nucleotide changes were observed among the Dutch outbreak sequences, whereas the 1,621 aa RdRp protein, which contains the small replicase subunit, showed 100% to 99.8% similarity on protein level (up to 6 aa changes). The 267 aa movement protein displays relatively the highest level of diversity on protein level with 100% to 98.1% aa similarity (up to 5 aa changes).

Bayesian inference of phylogeny results in clustering of the ToBRFV genomes sequenced from the phytosanitary inspections of greenhouses growing tomato in the Netherlands in three main clusters (Fig 2: red, blue and green clusters). A fourth cluster was formed with sequences obtained from tomato fruits from one lot intercepted during import from Egypt (Fig 2: orange cluster). Clustering obtained with RAxML, which is part of the Nextstrain build, results in the same grouping as the Bayesian analysis for the viral strains obtained from the Dutch outbreak locations, but some rearrangements can be seen between clusters (S3 Fig).

**Figure 2.**
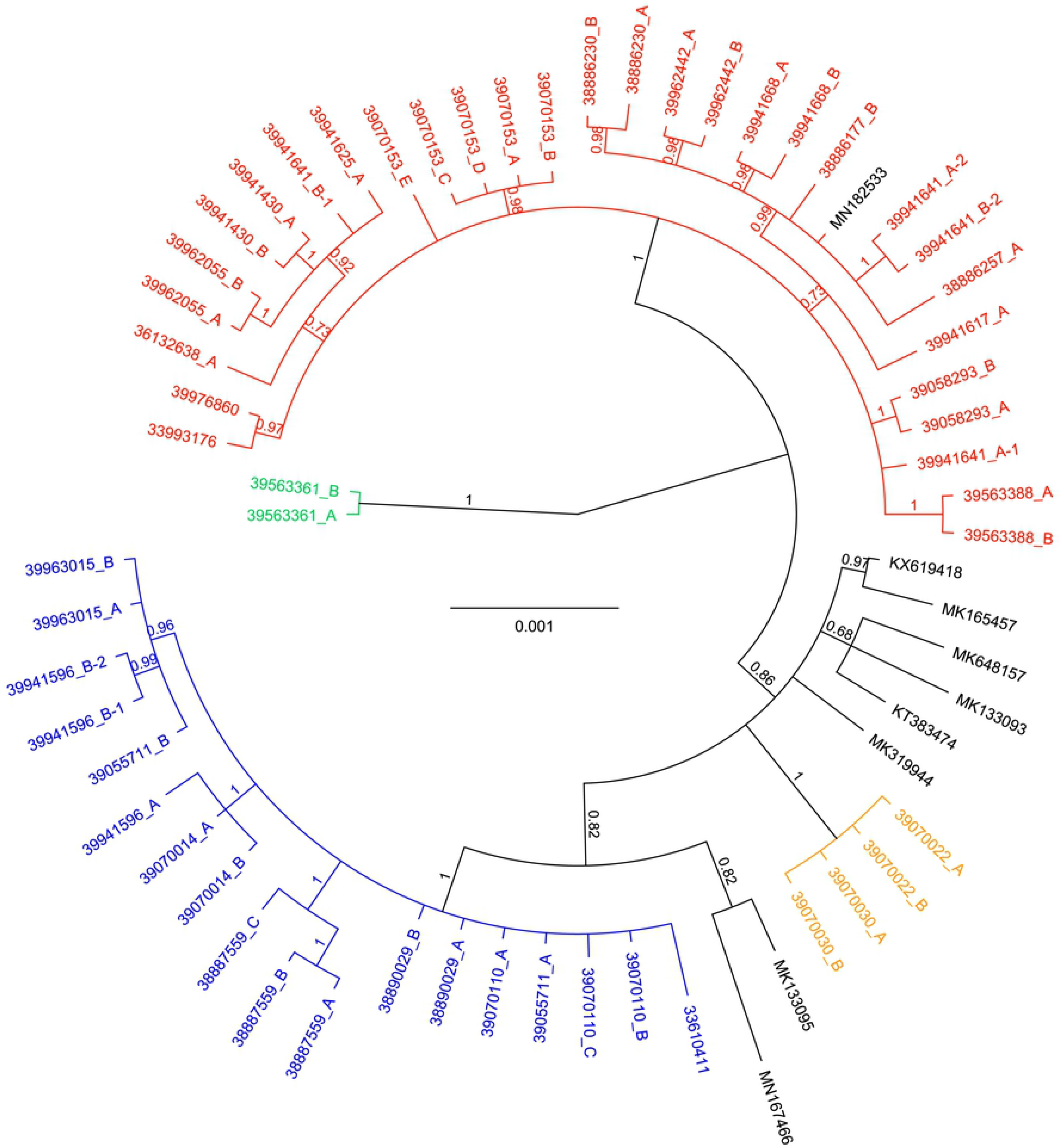
Bayesian inference of phylogeny based on a 6,404 nt alignment of 63 complete ToBRFV genomes. Viral strains from the Dutch outbreak are found in three separate clades which have been colored red, blue and green. Four ToBRFV genome sequences generated from material intercepted during an import inspection from Egypt form a separate cluster (orange). Bayesian posterior probabilities (PP) are displayed at the branch nodes. The scale bar indicates the number of substitutions per site.

In twenty datasets, one dominant ToBRFV genotype was detected (major genotype ≥98%), but in the remaining 34 datasets intermediate SNPs were observed suggesting the presence of multiple genotypes within the sample (Fig 3). Within sample diversity of major genotypes ranged from 54.5% to 89.0% in the isolates with mixed genotypes.

**Figure 3.**
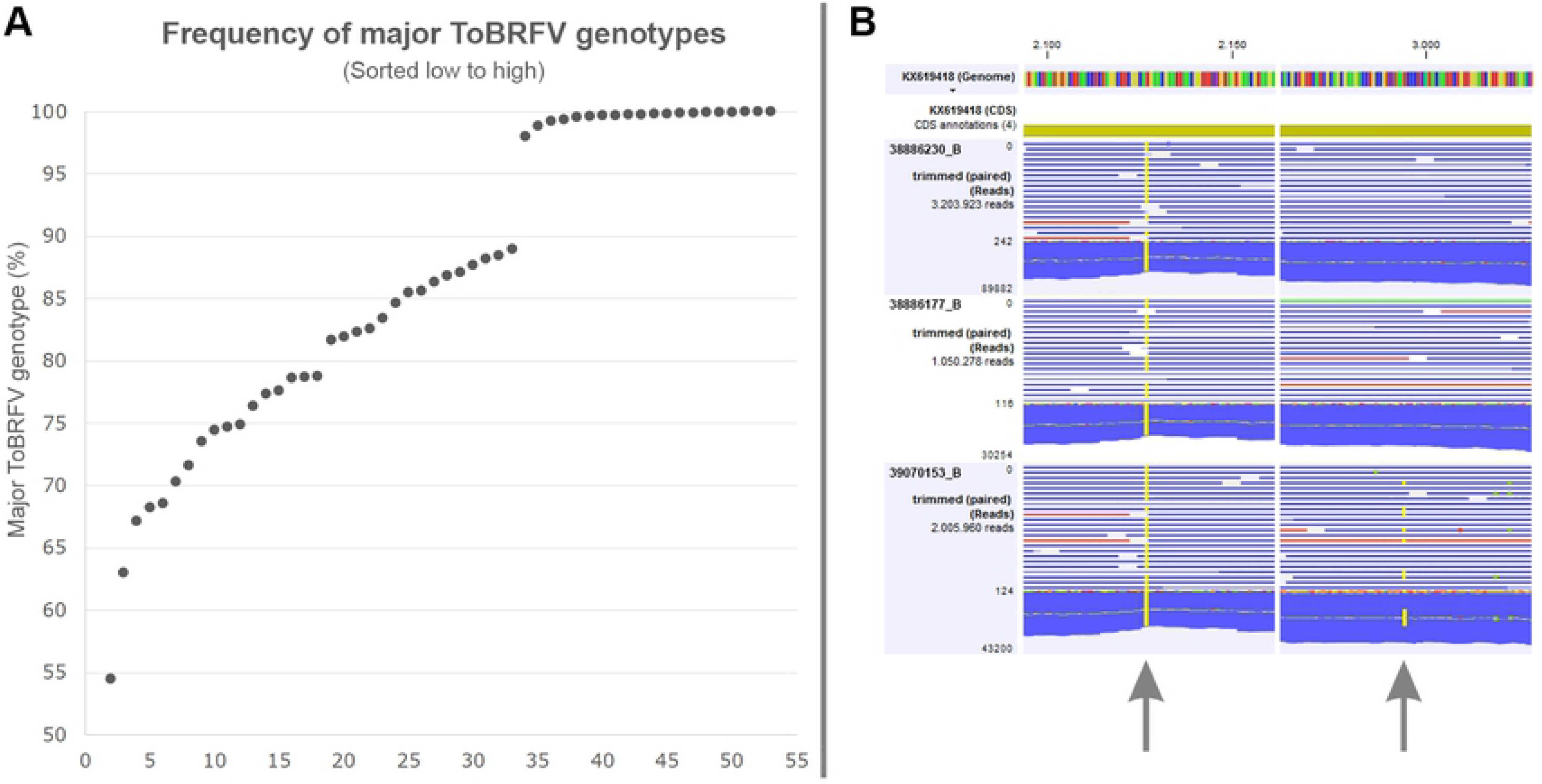
Within sample diversity of ToBRFV genotypes. **A.** Frequency (y-axis) of major ToBRFV genotypes in datasets (x-axis) generated in this study. Within sample frequencies of major genotypes are sorted low to high. **B.** Detail of RNAseq read mappings to the small replicase subunit of KX619418 for ToBRFV samples 38886230_B, 38886177_B, and 39070153_B. For each sample the total number of mapped reads is shown as well as the maximum read coverage. The left SNP (arrow) represents a single genotype (>99.7%) that is present in all three samples. The right SNP reveals the presence of a minor genotype (32.8%) in sample 39070153_B.

### Nextstrain build

The ToBRFV Nextstrain build contains 63 complete ToBRFV genomes generated from material sampled between October 2014 and November 2019. Information in the build is presented in three main panels: clustering of genomic diversity, geographical origin of the samples, and diversity relative to the ToBRFV Tom1-Jo genome (KT383474 = NC_028478) (Fig 4).

**Figure 4.**
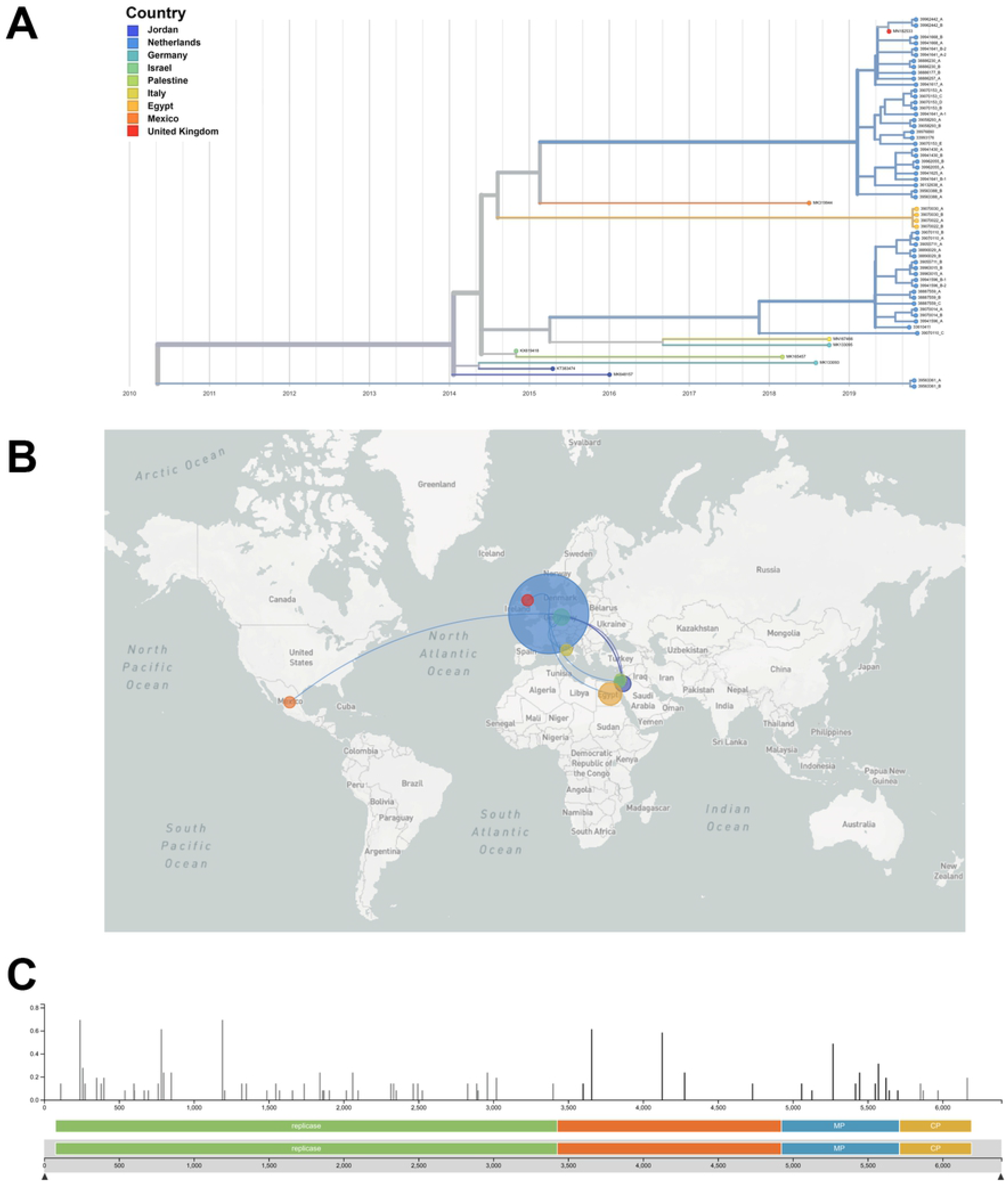
Genomic epidemiology of ToBRFV from October 2014 to November 2019. The Nextstrain build consists of three main linked panels: a phylogenetic tree (**A**), geographical distribution and most likely distribution events (**B**), and genomic diversity represented in the dataset using the ToBRFV Tom1-Jo genome (KT383474 = NC_028478) as reference with the small replicase subunit annotated in green, the RNA-dependent RNA polymerase in orange, the movement protein in blue and the coat protein in yellow.

The associated metadata included in the build allows users to color the nodes in the tree according to the host plant, and the (anonymized) rootstock and scion varieties from which the sequence was obtained. In addition, the within-sample genomic diversity is a trait that can be visualized. Internal node colors indicate the predicted ancestral state of a given trait, and the confidence of that state is conveyed by saturation of the color of the internal node. The cladogram can be shown in different styles such as rectangular, radial and unrooted. The branch lengths of the tree can be shown based on divergence or in function of time. Based on the information provided in the build, Nextstrain determines the most likely transmission events, which can be animated from the webpage. The genotypes represented in the tree are plotted on a map, and users can set different levels of geographical resolution, i.e. continent, country, state and municipality (when this information is available). This allows simultaneous interrogation of phylogenetic and geographic relationships, with additional relevant metadata. Users can download metadata and tree-files from the webpage, and can create screenshots of their views.

## Discussion

In October 2019, the Dutch NPPO received a tomato sample that was suspected to be infected with ToBRFV. The sample formed part of an official specific survey of tomato fruit growers in the Netherlands carried out in the period July – October 2019. During the official survey in total 124 companies were visited and inspected for possible symptoms of ToBRFV. In total 47 samples were tested, whereby one sample tested positive. Following initial testing with ELISA, indicator plants and real-time RT-PCR, presence of ToBRFV was verified with analysis of plant rRNA depleted Illumina RNA sequence data. Since little was known about the intraspecific variation and the validation status of the diagnostic tests, it was decided to use RNAseq analysis for ToBRFV verification in the follow-up survey and intensified specific surveillance by the NPPO-NL.

Several primer pairs have been described for conventional RT-PCR that can be Sanger sequenced for ToBRFV verification. As a result, some authors have sequenced either the 5’ end of the small replicase subunit, the 3’ end of the small replicase subunit, the 3’ end of RdRp, the partial MP and CP, or a combination of these loci (S4 Fig). This inevitably results in incomplete datasets. In contrast to Sanger sequencing (33), RNAseq analysis with plant rRNA depleted Illumina data does not rely on the specificity of primers used to amplify the partial virus genome. Furthermore, instead of sequencing only a small part of the genome, Illumina RNAseq data allows the *de novo* assembly of the (near) complete ToBRFV genome. With the complete genome, all genetic information can be used to determine potential linkages between genotypes and epidemiological traits.

The real-time RT-PCRs used for the detection of ToBRFV have been validated for detection of the virus in tomato and pepper seeds (available on request from Naktuinbouw). When implementing the test for tomato leaves, in some cases late Cq values were obtained with the healthy tomato matrix (data not shown). To compensate for these late Cq values, the Cq cut-off value was lowered from 32 to 30. Samples with Cq values lower than 25 could be verified with Illumina RNAseq data. When Cq-values between 25 and 30 were obtained, ToBRFV could be verified in a single case only. Seventy percent of the samples taken resulted in Cq values lower than 25. When ToBRFV suspected samples could not be verified, companies were revisited and additional samples were taken. Overall there is a good correlation between the Cq values and normalized read coverage in the samples analyzed (*R^2^*_CP assay_ = 85.3% and *R^2^*_MP assay_ = 76.4%; S5 Fig).

Visual inspection of the read mappings revealed that multiple genotypes existed within some samples. Only in two subsamples the variation between the genotypes within a single subsample was high enough to produce two separate contigs from the *de novo* assembly. In other cases, the minor genotypes were not manually reconstructed from the assemblies as they could represent one or more than one minor genotypes. When looking at the within sample diversity, a gap of almost 10% between samples with one clear major genotype and samples with mixed genotypes can be seen (Fig 3). Since reads were mapped to reference KX619418, variants mapping between 89.0 and 98.0 % would have been detected if they were present. ToBRFV populations with only a single major genotype could have undergone a recent genetic bottleneck event. However, no correlation between within sample genomic diversity and genotype or any of epidemiological traits analyzed was observed.

### Epidemiology of Dutch ToBRFV outbreak

The ToBRFV Nextstrain build contains 63 complete ToBRFV genomes, 54 of which have been generated in this study. The genomic sequences generated from the Dutch outbreak sites group in three main clusters (Fig 2: red, blue, and green clusters). The TimeTree analysis estimates the divergence of the red and blue clusters well before the first finding of ToBRFV in the Netherlands (inferred date = May 2014, confidence interval = March 1980 – October 2014), and the three clusters are hypothesized to represent three different sources. The blue and green clusters consist of sequences that so far were only found in the Netherlands. One sequence obtained from an outbreak in the United Kingdom groups in the red cluster, suggesting that these are potentially linked. The genomes sequences from the Dutch outbreak locations display a broad range of genomic diversity. However, genotypes found at outbreak locations in the Middle East, North-America, Italy and Germany were not found in Dutch greenhouses. Furthermore, a genotype was found in a sample obtained from an import inspection from Egypt which had not been found before at any of the outbreak locations. Diversity is expected to be highest at the center of origin, and increased sampling at outbreak locations and presumed origin will aid further understanding of ToBRFV diversity. This will eventually allow the determination of the origin for this viral species.

In all but one of the inspected companies, ToBRFV genotypes obtained from the samples taken belonged to one of the main clusters. In a single company, ToBRFV genotypes from both the red and blue cluster were obtained. These two different genotypes were obtained from separate greenhouses of the same company, and it is likely that they were introduced independently.

A number of epidemiological traits were analyzed in context of genomic diversity to identify the possible source(s) of the ToBRFV outbreak in the Netherlands. These included rootstock and scion varieties, young plant providers, seed batches and seed providers. However, no linkages were identified between these epidemiological traits and companies or genotypes. Seen the distribution of epidemiological traits among genotypes and outbreak locations, it is likely that the virus has been present in the Netherlands some time before the first official samples were obtained and tested. Also, establishing the origin of Tobamoviruses could prove to be too challenging as they are easily transmitted and can spread rapidly over short periods in time when they remain unnoticed. Companies that have been found infected must take strict hygiene measures to prevent spread of the virus to other companies and to eradicate the virus at the end of the cropping season.

The ToBRFV Nextstrain build currently holds ToBRFV genomes from various outbreak locations in the Middle East, North-America and Europe. However, there is a strong bias towards genotypes obtained from the Dutch outbreak locations. Sampling bias and lack of data can hamper the determination of reliable transmission links. The predictive power of the tool will improve with the addition of additional genomes. Therefore, we encourage other organizations to share data or biological materials together with relevant metadata in order to improve the build. Such a community effort will contribute to source attribution and to a better understanding of the diversity and spread of ToBRFV worldwide. The ToBRFV Nextstrain build will be maintained by NPPO-NL.

## Acknowledgements

Many thanks to the NPPO-NL staff members Marieke van Lent, Leontine Colon, Bram Lokker, Isabel Stagg, and Stephanie Rensen, Lucas van der Gouw and Eveline Metz for collecting metadata, running tests and providing technical support with the generation and analysis of NGS data. The support by Naktuinbouw staff members Agata Jodlowska, André van Vliet, and Ruud Barnhoorn in performing the first-line detection tests is greatly acknowledged. Thank you Annelien Roenhorst and Martijn Schenk (NPPO-NL) for critically reviewing the manuscript before submission.

## Supporting information

**S1 Fig. Visualization of the CLC Genomics Workbench pipeline used to identify putative ToBRFV contigs.** The pipeline combines a quality trim for input reads, *de novo* assembly, blast based detection and visualization of blast output in Krona. The pipeline runs on all input data, and on a random sample of 1% of all reads. Blue process steps indicate output, and custom plugins created for the pipeline are named CEA (CLC external application).

**S2 Fig. Example of Krona visualization of blast output for sample 38886230_A.** When using all data for the *de novo* assembly, and after blast-based filtering of plant contigs, 1266 contigs remained. The majority of these sequences (99%) did not produce a blast hit, but 3 viral blast hits were obtained (left panel). When selecting the viral hits, the blast-based identity is shown (right panel). In this example, one ToBRFV hit and two PepMV hits are obtained. The sequence producing the ToBRFV hit is obtained via a link, which is then further analyzed to determine the presence of the virus in the sample.

**S3 Fig. Comparison of the Nextstrain RAxML tree (left) and the Bayesian inference of phylogeny (right) of 63 ToBRFV genomes included in this study.** Links are drawn for the isolates generated in this study. The Nextstrain tree is colored based on the country of origin, whereas the Bayesian tree is colored based on the main clusters identified for the viral genomes sequenced in this study. Grouping within the main clusters is identical between the two analyses, but some groups are placed at different positions in the overall phylogeny (e.g. the Egyptian sequences: orange cluster).

**S4 Fig. Publicly available ToBRFV sequences mapped to ToBRFV genome NC_028478 (= KT383474).** Coding sequences are annotated in yellow, and sequences identical to the reference sequence are shown in grey. Differences relative to the reference are highlighted in black. Apart from the nine complete genome sequences, 37 Sanger sequences have been submitted to NCBI covering ① 5’ end of the small replicase subunit, ③ 3’ end of the small replicase subunit, ④ 3’ end of RdRp, and ⑤ the partial MP and CP.

**S5 Fig. Correlation between Cq values and read normalized read mapping.** Real-time RT-PCR Cq values of the CP (top) and MP (bottom) assays are displayed on the x-axis and the normalized average read-coverage is shown on the y-axis. To allow comparison between the different datasets, the average read coverage of the ToBRFV genome for a given sample was multiplied by the fraction of reads generated for that sample relative to the mean of reads generated for all samples.

**S1 Table. Details of real-time RT-PCR detection and RNAseq-based verification tests performed on ToBRFV samples included in this study.**

